# Ergosterol distribution controls surface structure formation and fungal pathogenicity

**DOI:** 10.1101/2023.02.17.528979

**Authors:** Hau Lam Choy, Elizabeth A. Gaylord, Tamara L. Doering

## Abstract

Ergosterol, the major sterol in fungal membranes, is critical for defining membrane fluidity and regulating cellular processes. Although ergosterol synthesis has been well defined in model yeast, little is known about sterol organization in the context of fungal pathogenesis. We identified a retrograde sterol transporter, Ysp2, in the opportunistic fungal pathogen *Cryptococcus neoformans*. We found that the lack of Ysp2 under host-mimicking conditions leads to abnormal accumulation of ergosterol at the plasma membrane, invagination of the plasma membrane, and malformation of the cell wall, which can be functionally rescued by inhibiting ergosterol synthesis with the antifungal drug fluconazole. We also observed that cells lacking Ysp2 mislocalize the cell surface protein Pma1 and have thinner and more permeable capsules. As a result of perturbed ergosterol distribution and its consequences, *ysp2*Δ cells cannot survive in physiologically-relevant environments such as host phagocytes and are dramatically attenuated in virulence. These findings expand our knowledge of cryptococcal biology and underscore the importance of sterol homeostasis in fungal pathogenesis.

**IMPORTANCE:** *Cryptococcus neoformans* is an opportunistic fungal pathogen that kills over 100,000 people worldwide each year. Only three drugs are available to treat cryptococcosis, and these are variously limited by toxicity, availability, cost, and resistance. Ergosterol is the most abundant sterol in fungi and a key component in modulating membrane behavior. Two of the drugs used for cryptococcal infection, amphotericin B and fluconazole, target this lipid and its synthesis, highlighting its importance as a therapeutic target. We discovered a cryptococcal ergosterol transporter, Ysp2, and demonstrated its key roles in multiple aspects of cryptococcal biology and pathogenesis. These studies demonstrate the role of ergosterol homeostasis in *C. neoformans* virulence, deepen our understanding of a pathway with proven therapeutic importance, and open a new area of study.

## INTRODUCTION

*Cryptococcus neoformans* is a fungal pathogen that causes 112,000 HIV-associated deaths per year and accounts for 19% of AIDS-related mortality (1). During infection, spores or desiccated yeast cells are inhaled, resulting in pulmonary infection. In immunocompetent hosts, cryptococcal infections are generally asymptomatic and are either cleared or remain latent. However, in immunocompromised patients, the fungi disseminate from the lungs and enter the central nervous system, resulting in often-fatal meningoencephalitis (2–4).

Ergosterol is the most abundant sterol in fungal membranes (5). It is critical in defining membrane fluidity and permeability and regulating protein sorting and the activity of membrane-associated enzymes (5, 6). Beyond basic biology, ergosterol is also an important therapeutic target for *C. neoformans* infections. Treatment options for cryptococcosis are limited to three drugs: amphotericin B (AmB), fluconazole, and flucytosine (7, 8). Of these, AmB and fluconazole target ergosterol itself or its biosynthetic pathway (9–11). Ergosterol synthesis has been well-defined in model yeast *Saccharomyces cerevisiae* (6, 12), but less is known about sterol organization, particularly in the context of fungal pathogenesis.

Although sterols are synthesized in the endoplasmic reticulum (ER), most are transported to other organelles, notably the plasma membrane (PM), which contains up to 90% of cellular sterols (13, 14). This movement is mainly mediated by sterol-specific lipid transport proteins, which are independent of the secretory pathway (15). *S. cerevisiae* expresses two families of these proteins: oxysterol-binding proteins (OSH) and lipid transfer proteins anchored at membrane contact sites (LAM). The seven cytosolic OSH proteins localize at membrane contact sites and move sterols to and from the ER in exchange for other lipids (16–18). The recently identified LAM proteins are anchored by transmembrane domains and possess characteristic StART (Steroidogenic Acute Regulatory Transfer)-like domains that bind sterols (Fig 1A) (19). In *S. cerevisiae*, this family is represented by six proteins, Lam1-Lam6 (19). Lam1-4 are localized at sites of ER-PM membrane contact, while Lam5 and Lam6 are localized at sites of contact between the ER and mitochondria or vacuoles (19, 20). Lam2, also called Ysp2, has been suggested to be a retrograde sterol transporter that moves sterols from the PM to the ER and may be important for mitochondrial morphology (19, 21).

**Fig 1.**
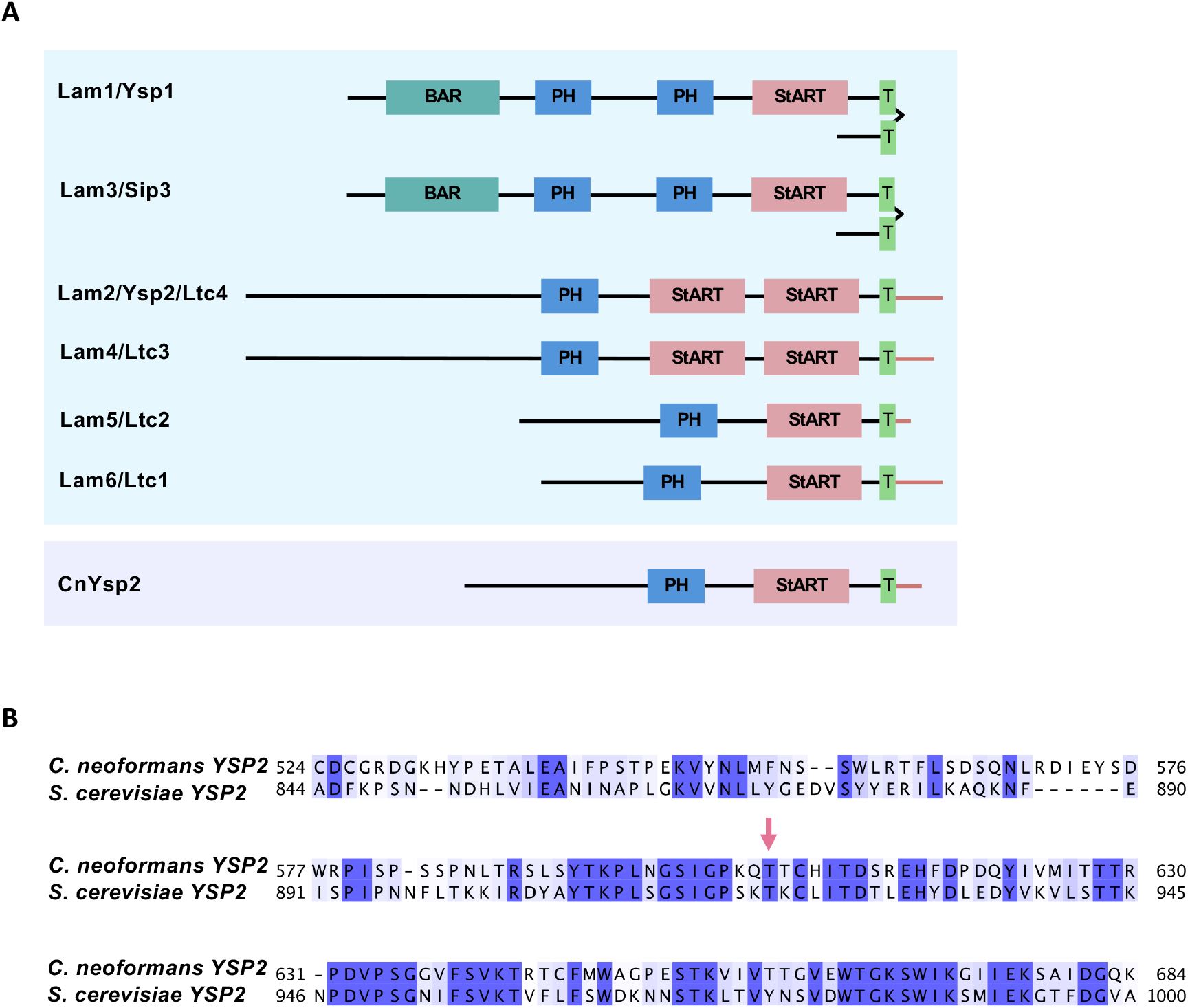
Alignment of cryptococcal Ysp2 with *S. cerevisiae* homologs. (A) LAM family proteins in *S. cerevisiae* (blue box) and *C. neoformans* (purple box). Domain abbreviations are Bar, Bin/amphiphysin/RVS; PH, pleckstrin-homology; StART, Steroidogenic Acute Regulatory Transfer-like; and T, transmembrane. (B) Alignment of *C. neoformans* and *S. cerevisiae YSP2* StART-like domains, with CLUSTALX coloring of conserved residues. Pink arrow, residue predicted to make van der Waals contact with ergosterol (25).

Mechanisms of sterol transport have been examined in model yeast but not in fungal pathogens, and the relationship between sterol organization and fungal pathogenesis remains unexplored. To tackle this question, we investigated the role of the only apparent LAM family member in *C. neoformans*. This protein, named Ysp2 for its homology to the *S. cerevisiae* transporter, was previously identified in a caspofungin sensitivity screen and shown to influence membrane integrity (22). We found that lack of Ysp2 under conditions that mimic the mammalian host environment leads to excess accumulation of ergosterol at the PM, invagination of the PM, and striking malformation of the cell wall. These processes can be functionally rescued by inhibiting ergosterol synthesis with fluconazole. We also observed perturbations of sterol synthesis and storage in *ysp2*Δ mutant cells. We conclude that Ysp2 is a retrograde sterol transporter that is critical for survival in host environments and cryptococcal virulence.

## RESULTS

### Ysp2 is required for *in vivo* and *in vitro* virulence

We used the *S. cerevisiae* Ysp2 protein sequence to identify the homologous cryptococcal gene, *CNAG_00650*. BLASTp searches of the *S. cerevisiae* genome using this sequence yield the original *S. cerevisiae* gene; based on this reciprocity the *C. neoformans* protein has the same name. Ysp2 in *C. neoformans* and *S. cerevisiae* have 39% amino acid identity overall, with an E value of 6 × 10^−51^ (Fig 1B shows homology in the StART-like domain). For functional studies, we generated a *ysp2Δ* deletion mutant in *C. neoformans* strain KN99α (referred to as WT below), and also complemented it at the native locus (referred to as *YSP2*).

To investigate the role of Ysp2 in fungal pathogenesis, we assessed the virulence of *ysp2*Δ in a mouse model of cryptococcosis, where disease progression is monitored by weight loss. All mice infected with WT or complemented strains steadily lost weight (Fig S1A) and succumbed to infection by day 22, but mice infected with *ysp2*Δ showed no signs of illness and were only sacrificed when the experiment was terminated at day 80 (Fig 2A). Consistent with these findings, the lungs and brains of mice infected with WT or complemented strains showed high fungal burden at sacrifice, while those of mice infected with *ysp2*Δ yielded minimal fungi (Fig 2B).

**Fig 2.**
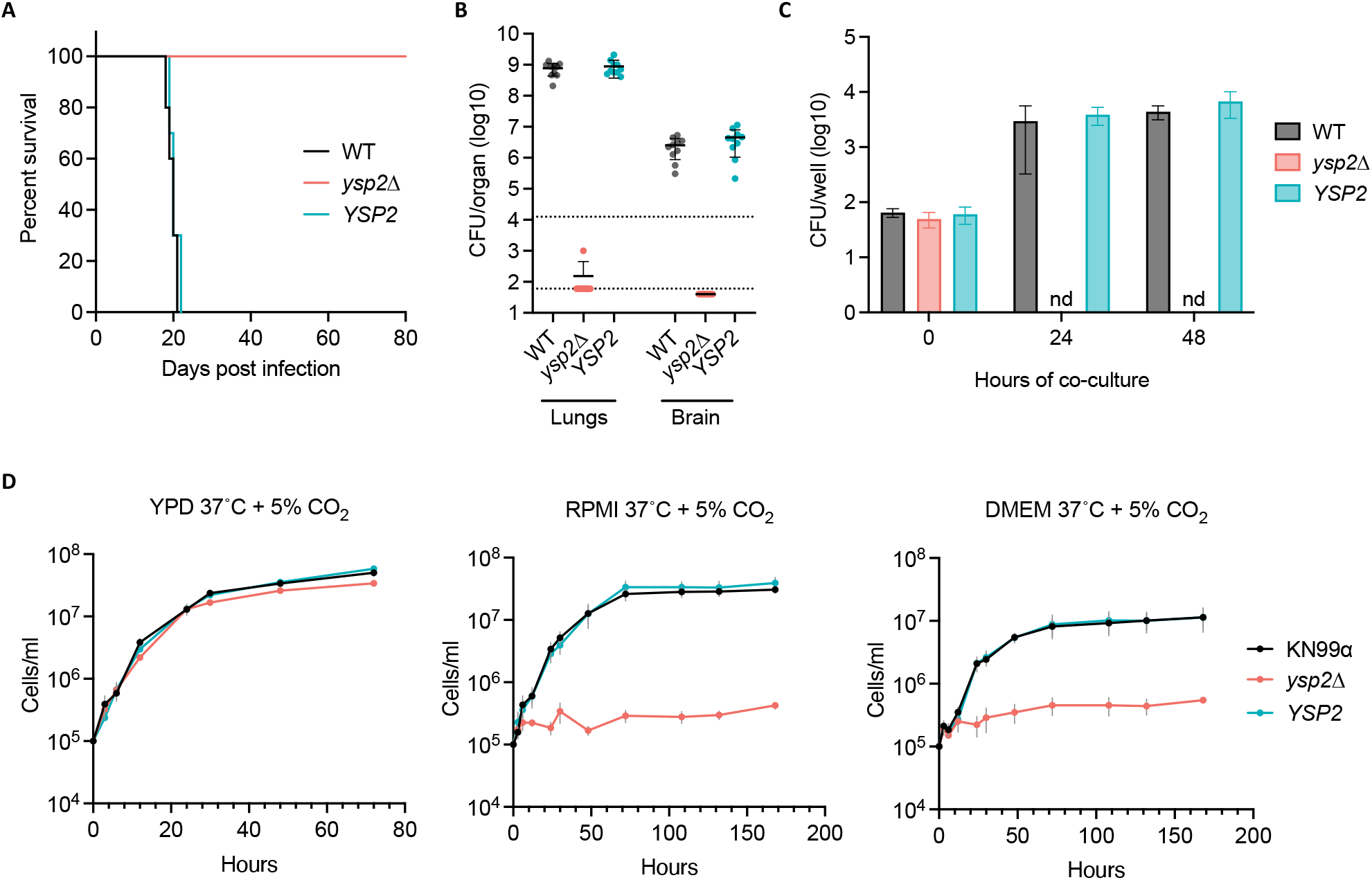
Ysp2 is required for virulence and survival in physiological environments. (A) Survival of C57BL/6 mice after intranasal infection with 1.25 × 10^4^ fungal cells, with sacrifice triggered by weight below 80% of initial weight. (B) Lung and brain fungal burdens of mice from Panel A at sacrifice. Top dotted line, initial inoculum; bottom dotted line, limit of detection. (C) *In vitro* survival of cryptococci. BMDMs were co-incubated with the indicated strains (1.5 h, MOI = 0.1) and washed to remove free fungi before lysis at the times shown and assessment of cryptococcal CFU. nd, not detected. Mean ± SD is plotted; results shown are representative of at least two biological replicate experiments. (D) Growth curves in the conditions shown (mean ± SEM of three independent experiments).

*C. neoformans* is a facultative intracellular pathogen, which may enter and survive within host phagocytes (23, 24). To determine the effect of Ysp2 on these processes, we examined fungal interactions with host macrophages. We found that *ysp2*Δ cells were phagocytosed at the same rate as the WT and complemented strains (Fig 2C and S1B) but were much more susceptible to killing after internalization; they were completely cleared by 24 hours of incubation, while the control populations significantly increased in that interval (Fig 2C).

We wondered whether the severe attenuation of *ysp2*Δ cells in mice and in macrophages reflected susceptibility to features of the host environment, independent of specific host responses. Upon testing this, we found that mutant growth in rich medium was not perturbed by exposure to host physiological temperature (37°C) or CO_2_ level (5%) (Fig 2D and S2). However, the population no longer increased when the conditions were changed to incorporate mammalian tissue culture medium along with these environmental changes (Fig 2D). Notably, the *ysp2*Δ cells remained viable under these conditions at all times shown, as demonstrated by their ability to form colonies upon transfer to rich medium (Fig S2C).

### *C. neoformans* lacking Ysp2 exhibits defects in surface structures

To explore factors that might influence the pathogenicity of *C. neoformans* lacking Ysp2, we first examined the best-known cryptococcal virulence factor, its polysaccharide capsule. This structure presents a physical barrier to phagocytosis and modulates the host immune response (24–26). For cells grown in host-like conditions, the capsule thickness of *ysp2*Δ was reduced by almost 40% compared to control strains (Fig 3A). Interestingly, despite their thinner capsules, the mutant cells bound more anti-capsule antibodies (Fig 3B). These apparently contradictory results suggested a possible change in the mutant capsule architecture. We tested this idea by measuring the permeability of capsule to 2,000 kDa fluorescent dextran beads, using cell wall staining with calcofluor white (CFW) to mark its inner boundary. The beads penetrated twice as deeply into the capsule of *ysp2*Δ cells compared to WT (Fig S3A), supporting our hypothesis. These studies also revealed some intriguing deformities of the mutant cell wall (Fig. 3B, CFW staining), which are pursued below.

**Fig 3.**
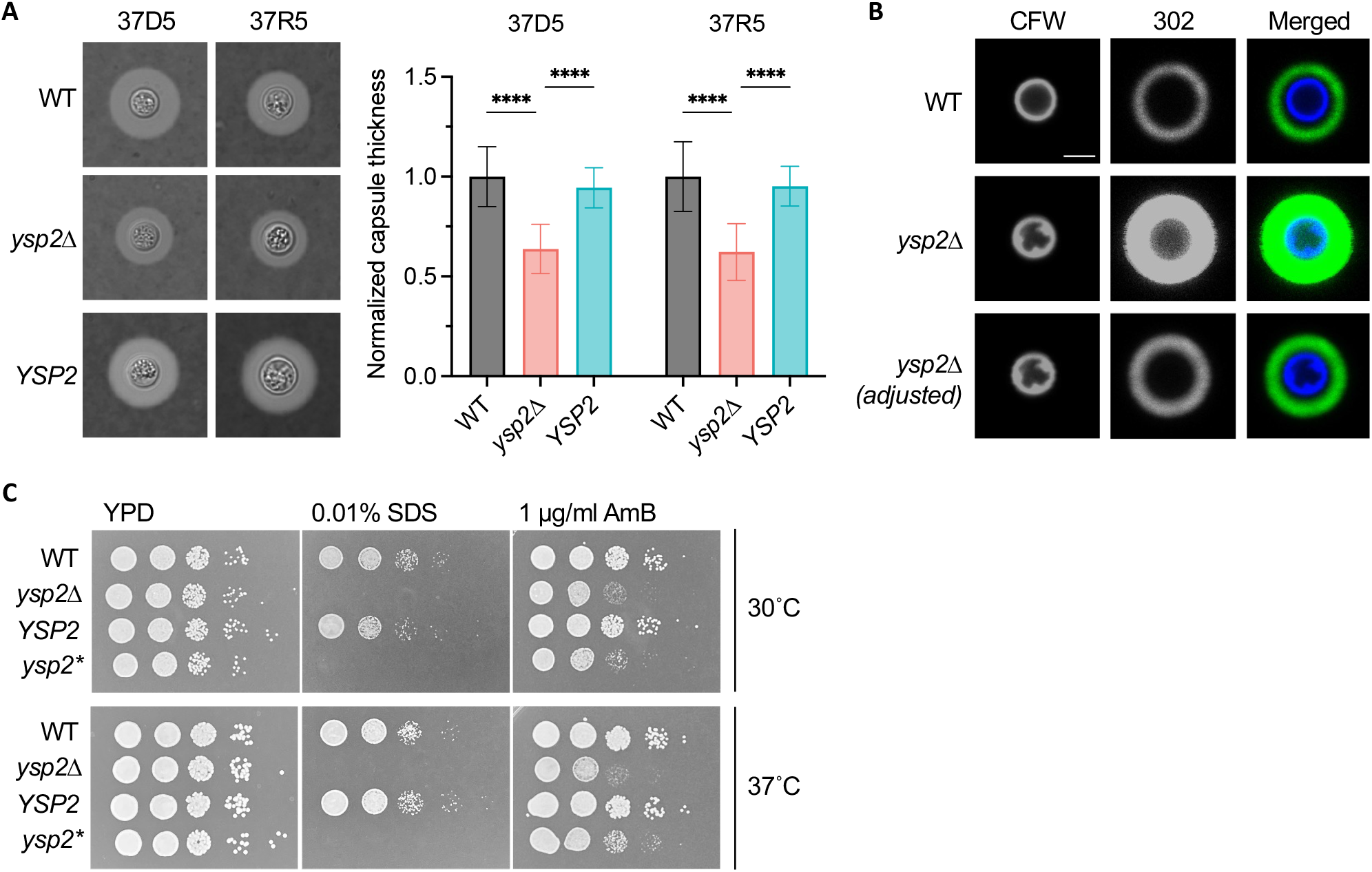
The *ysp2*Δ mutant exhibits cell surface defects. (A) Capsule thickness. The indicated strains were grown in host-like conditions (37°C, 5% CO2, with either DMEM (37D5) or RPMI (37R5)), stained with India ink (left), and capsule thickness was measured with ImageJ and normalized to cell radius and WT value (right). Mean ± SD of at least 50 cells per sample are shown. ****, P<0.0001 by one-way ANOVA. (B) Representative confocal micrographs of the indicated strains after growth in 37R5 for 24 h and staining with CFW (cell wall) and MAb 302 conjugated to Alexa Fluor 488 (capsule). Images in the first and second rows were obtained at the same gain and intensity settings; in the third row, confocal gain of the *α*-capsule Ab channel was reduced. All images are to the same scale; bar, 5 μm. (C) Serial 10-fold dilutions of the indicated strains in the conditions shown.

If Ysp2 is indeed a retrograde sterol transporter, its absence may alter lipid distribution and thereby compromise membrane integrity. To test this hypothesis, we plated serial dilutions of *ysp2*Δ cells in the presence of membrane stressors. Compared to controls, *ysp2*Δ was far more sensitive to SDS (Fig 3C). It was also more sensitive than control strains to the ergosterol-binding antifungal compound amphotericin B (AmB) (Fig 3C), as was previously observed in *C. neoformans* (22) and in the corresponding *S. cerevisiae* mutant (19).

We suspected that the aberrant phenotypes of *ysp2*Δ were due to its inability to appropriately distribute sterols. In *S. cerevisiae*, residue T921 of Ysp2 is required for ergosterol binding and consequent retrograde transfer activity (27). We used Clustal Omega to identify T606 as the corresponding amino acid in *C. neoformans* and mutated the *YSP2* gene to replace it with aspartic acid (Fig 1B). The resulting strain, *ysp2*\*, phenocopied the deletion mutant *ysp2*Δ (Fig 3C), supporting our model that defective ergosterol binding causes the observed phenotypes and attenuated virulence.

Ysp2 has been implicated in mitochondrial morphology in model yeast (21), probably because of the critical roles sterols play in the biogenesis and maintenance of mitochondrial membranes (28, 29). When we stained cells with MitoTracker CMXRos to assess *C. neoformans* mitochondrial morphology, we saw bright staining that was absent in WT (Fig S3B). Because the accumulation of this compound depends on mitochondrial membrane potential, we wondered whether this characteristic was altered in *ysp2*Δ. When we stained the mutant with TMRE, a cationic dye that accumulates in mitochondrial inner membranes based on membrane potential, we observed a broad peak, with roughly 50% of cells exceeding control staining (Fig S3C). However, *ysp2*Δ cells exhibited no growth defects on media containing alternative carbon sources or electron transport chain-inhibiting compounds (Fig S3D). We conclude that despite its effects on mitochondrial membranes, Ysp2 has minimal impact on mitochondrial function.

Based on Ysp2’s putative role in retrograde transport of sterols from the PM, we next focused our attention on this structure. To examine surface morphology, we stained cells with filipin, a fluorescent dye which binds sterols, and the cell wall dye Lucifer Yellow. Compared to the smooth ring staining patterns of control strains, *ysp2*Δ cells grown in host-like conditions showed irregular invaginations in both filipin and Lucifer Yellow signal (Fig 4A), similar to what we had noted earlier with CFW (Fig. 3B). The mutant cells additionally showed brighter filipin fluorescence (Fig 4A and 4B); this suggested higher sterol levels, which would also be consistent with impaired sterol removal from the plasma membrane. For a more detailed view of this striking phenotype, we examined the cells using transmission electron microscopy. Corroborating our light microscopy results, WT cells displayed even curvature of the cell wall and underlying PM. In contrast, the mutant showed distorted areas of both structures, which were only present when the cells were grown in host-like conditions (Fig 4C). Furthermore, in some regions, layers of wall material appeared to surround both membranous material and cytoplasmic content (Fig 4C).

**Fig 4.**
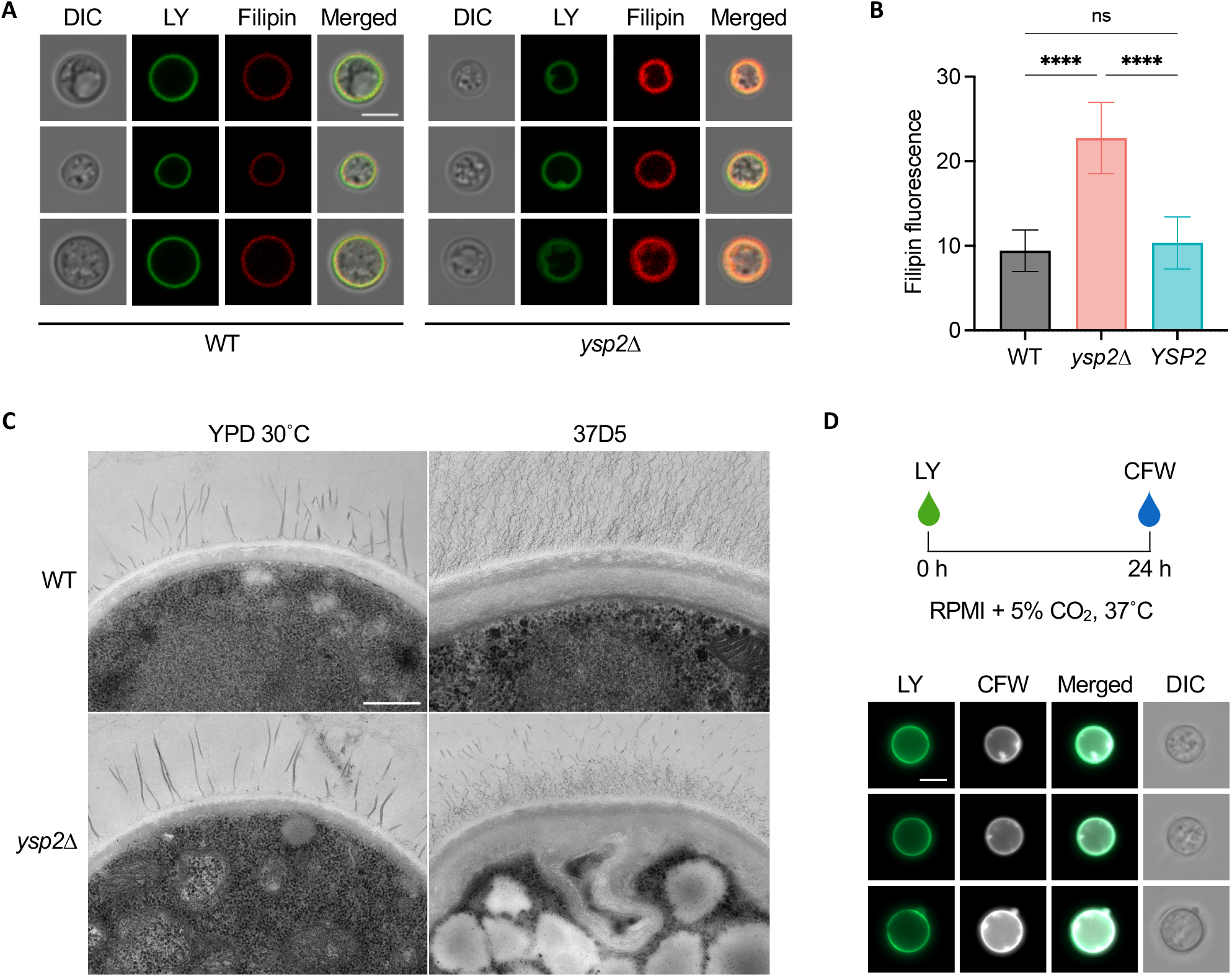
The *ysp2*Δ mutant exhibits malformations of the cell wall and plasma membrane. (A) Representative cells that were grown in 37R5 for 24 h and stained with Lucifer Yellow (LY) for cell wall and filipin for non-esterified sterols. All images are to the same scale; bar, 5 μm. (B) Filipin fluorescence (mean ± SD of mean grey value for at least 50 cells per sample). (C) Transmission electron micrographs of cells grown in 37D5. All images are to the same scale; bar, 500 nm. (D) Representative *ysp2*Δ cells that were stained with LY, grown in 37R5 for 24 h, and stained with CFW. All images are to the same scale; bar, 5 μm.

Intrigued by the unusual cell surface deformations of cells lacking Ysp2, we examined the kinetics of their formation. To do this, we grew cells in rich medium, stained their cell walls with Lucifer Yellow, cultured them in host-like conditions for 24 hours, and then stained them with CFW. In these studies, the Lucifer Yellow introduced before the culture period occurred as a smooth ring (Fig 4D) ; the surface invaginations were evident only in the CFW staining following 24 hours of growth. This suggests that the deformed regions are composed of newly synthesized cell wall material produced during the period of growth in host-like conditions, rather than being composed of older cell wall material that was somehow rearranged.

### Protein localization

Lipid rafts, or detergent-resistant microdomains, are ordered domains of the PM that are enriched in sphingolipids and sterols (30). An important function of these microdomains is to concentrate GPI-anchored polypeptides (30). In *C. neoformans* one of these is Pma1, a PM ATPase that is required for survival within host cells (31). To determine whether sterol accumulation in the PM would affect raft proteins, we generated strains expressing Pma1-mNeonGreen from the endogenous locus in both WT and *ysp2*Δ backgrounds. Although Pma1 normally localizes primarily to the PM in a uniform pattern, in *ysp2*Δ cells it also appeared as bright puncta (Fig 5A). This pattern occurred in roughly 70% of *ysp2*Δ cells, compared to 4% of WT (Fig 5A). This mislocalization of Pma1 in the mutant background suggests disruption of lipid rafts.

**Fig 5.**
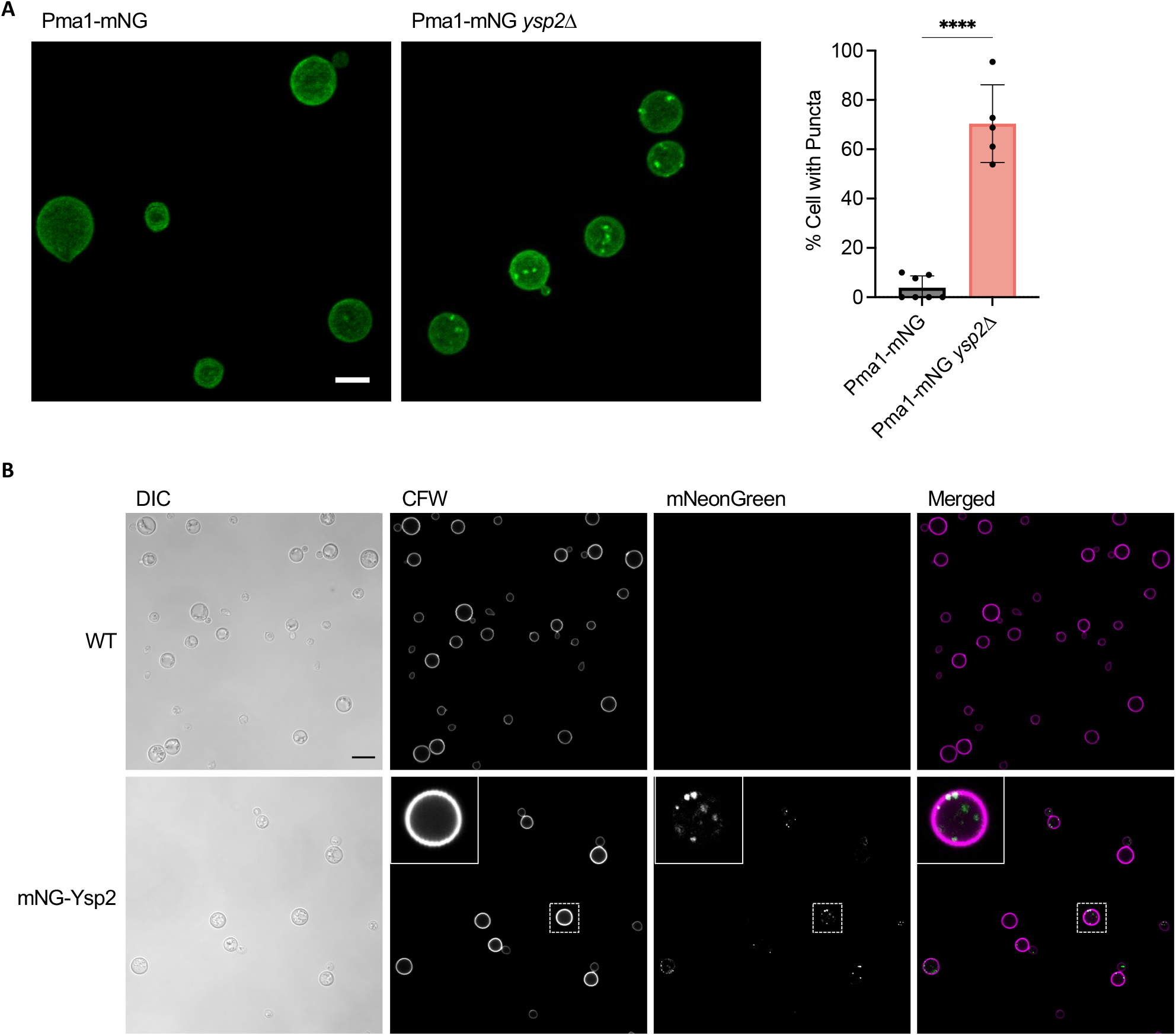
Protein localization imaged by confocal microscopy. (A) Pma1-mNeonGreen (Pma1-mNG) expressed in WT and *ysp2*Δ cells. Images, maximum intensity projections that sum Z-stacks. Both are to the same scale; bar, 5 μm. Plot, percent of cells with fluorescent puncta (mean ± SD based on at least 50 cells across 5 to 7 image fields). (B) Fluorescence and DIC images of WT cells alone (top) or the same cells expressing mNeonGreen-Ysp2 (mNG-Ysp2). Inset, an example cell (boxed) enlarged 3-fold. Bar, 5 µm.

We also assessed the localization of Ysp2 itself, by engineering cells to express mNeonGreen-Ysp2 from the native locus (Fig 5B). In addition to puncta of fluorescent protein at the cell periphery (compare to CFW staining), we observed roughly 35% (± SD of 10%) of the Ysp2 signal within the cell, quite different from its homolog in model yeast (see Discussion).

### Ysp2 modulates sterol distribution but not abundance

Our staining experiments suggested that cells lacking Ysp2 have abnormally high ergosterol at the PM. This could potentially be due to either an increase in total sterol levels or aberrant sterol distribution. To test the first possibility, we measured total ergosterol of WT, *ysp2*Δ, the complemented mutant, and *erg3*Δ, a biosynthetic mutant that produces extremely low levels of ergosterol (32). We found that the ergosterol content of *ysp2*Δ was comparable to that of the WT and complemented strains (Fig 6A), arguing against an increase in this compound and supporting our alternative hypothesis of altered sterol distribution. Perturbed lipid distribution is further supported by our observation of more lipid droplets in *ysp2*Δ than in WT and complemented strains (Fig 6C; see Discussion).

**Fig 6.**
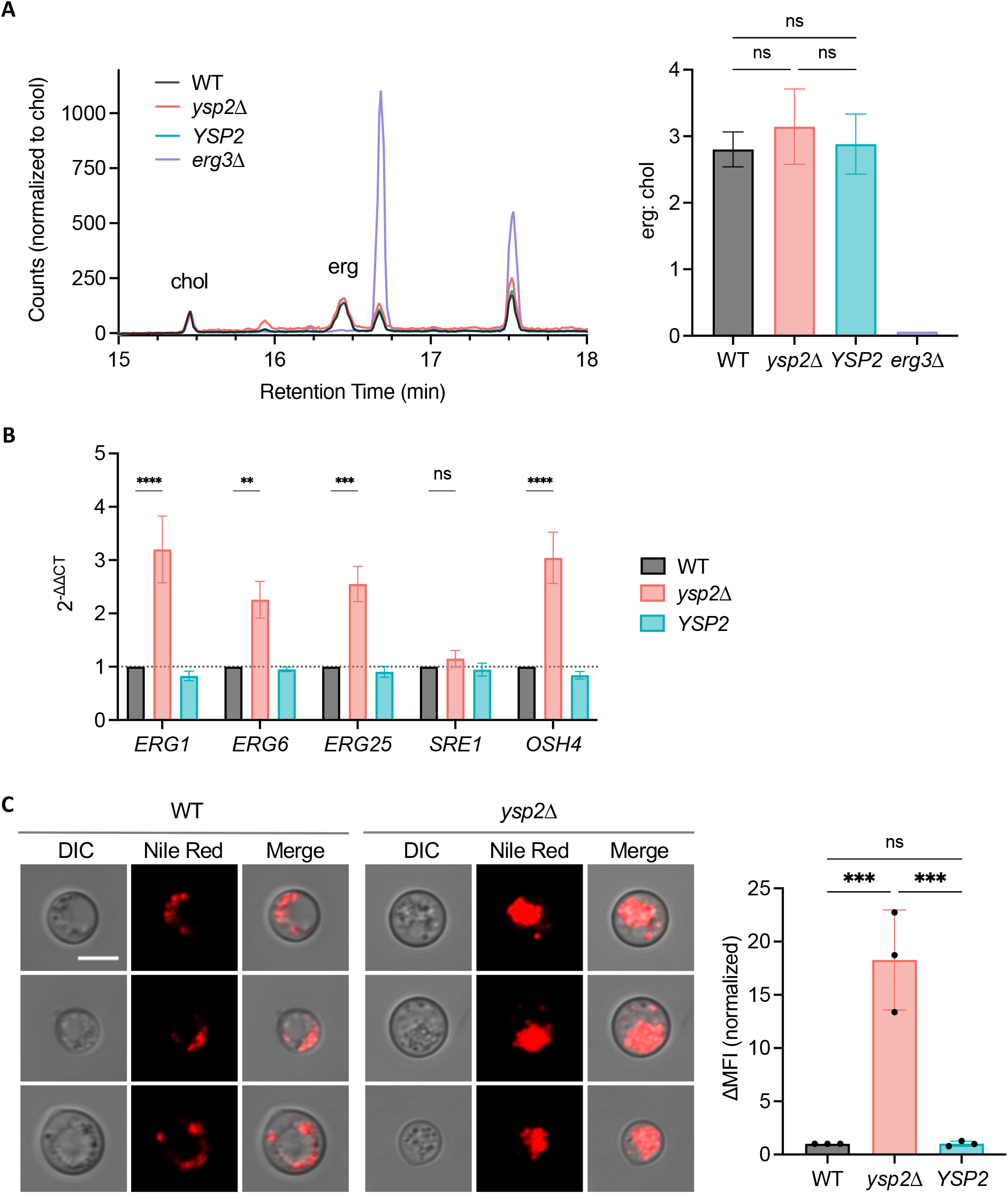
Sterol content, synthesis, and distribution. (A) Sterol content assessed by GC-MS. Left, a region of the total ion current chromatogram of total sterols from the indicated strains. Right, peak area of ergosterol (erg) normalized to cholesterol (chol, included as an internal standard). (B) Expression of ergosterol-related genes, measured by RT-qPCR and normalized to *ACT1* expression and WT values. The mean ± SEM of three independent experiments is shown. (C) Nile Red staining. Left, representative fluorescence images of WT and mutant strains. All images are to the same scale; bar, 5 μm. Right, ΔMFI from flow cytometry profiles of the indicated strains. Mean ± SD, normalized to WT, is shown for three independent experiments.

We also examined whether expression of sterol-related genes, chosen to represent various branches of sterol synthesis and transport, was altered in *ysp2*Δ cells. We detected modest upregulation of genes whose products act in ergosterol synthesis (*ERG1, ERG6*, and *ERG25* (6, 33)) and in transport of ergosterol away from its site of synthesis in the ER (*OSH4* (13, 28, 34, 35)) (Fig. 6B). However, because total ergosterol levels did not change, we speculate that any increased synthesis was likely balanced by sterol degradation or storage (see Discussion). The expression of *SRE1*, whose product regulates ergosterol synthesis but is itself regulated posttranscriptionally (36–39), was not affected (Fig. 6B).

### Drug sensitivity of *C. neoformans* depends on Ysp2

We previously observed that *ysp2*Δ cells are more sensitive than WT to the antifungal drug AmB (Fig 3C), which acts by binding and extracting ergosterol (10, 11). We hypothesized that this sensitivity was mediated by their increased cell surface ergosterol; consistent with this idea, these cells bound significantly more Cy5-conjugated AmB than control strains (Fig 7A and Fig S4B).

**Fig 7.**
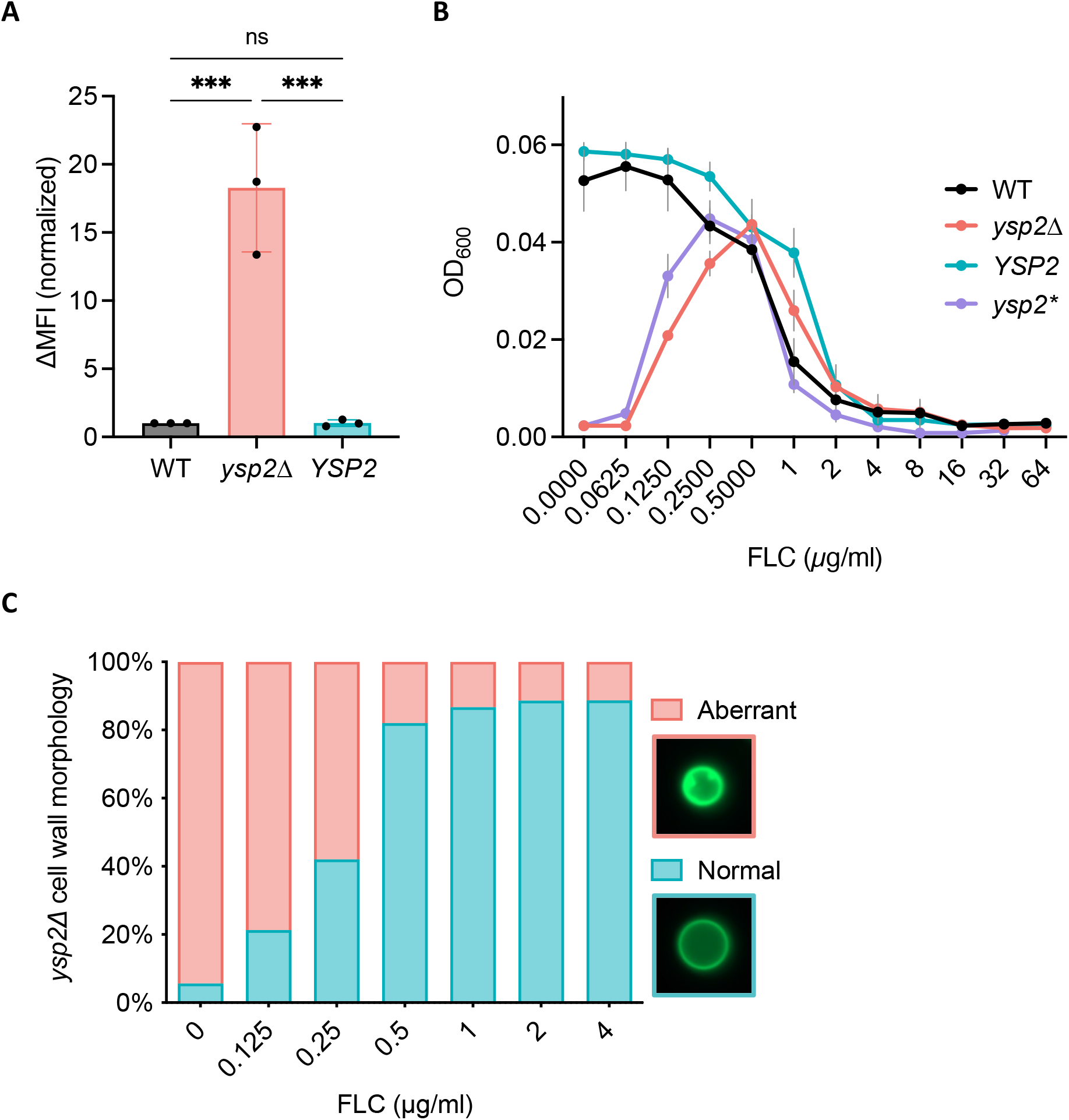
Antifungal drug interactions with *ysp2*Δ cells. (A) amphotericin B-Cy5 binding measured by flow cytometry. Mean ± SD of ΔMFI is shown for three independent experiments. (B) Mutant response to fluconazole (FLC). Cells were grown in 37R5 with FLC as indicated and OD600 measured at 48 h. Mean ± SEM of three independent experiments is shown. *ysp2\**, inactivated mutant. (C) Cells were grown in 37R5 and the fraction with aberrant cell walls was quantified at 24 h. 70 cells were scored per condition.

We further hypothesized that the excess PM ergosterol of the mutant cells led to the dramatic surface invaginations we had observed (Fig. 4). If true, the phenotype might be reversed by reducing sterol synthesis. To test this idea, we treated *ysp2*Δ cells with fluconazole, an antifungal that targets lanosterol demethylase (Erg11) and thereby inhibits ergosterol biosynthesis (9). Indeed, low levels of fluconazole showed a striking and dose-dependent rescue of *ysp2*Δ growth, such that it matched WT cell density at 0.5 *μ*g/ml (Fig 7B). This was accompanied by reversal of the cell wall invagination phenotype (Fig 7C) and plasma membrane irregularities (Fig S4C). Notably, *ysp2*Δ cells were not more resistant to fluconazole than WT, since at higher drug concentrations the growth of all strains was similarly reduced (Fig 7B). This contrasts with a previous study that showed *ysp2*Δ had slightly lower MIC for fluconazole (22), a difference we attribute to the distinct growth conditions used. We consistently observe more dramatic phenotypes (PM and cell wall morphologies) when we grow *ysp2*Δ cells are in tissue culture conditions versus rich medium, suggesting that fungal pathogens have a greater need for effective sterol organization in the host environment.

## DISCUSSION

Ergosterol, the major sterol of fungal membranes, is a key player in cell physiology and signal transduction and is also an important target of anti-cryptococcal drugs. Despite its importance, sterol organization in the context of pathogenesis has remained unexplored. In this study, we determined how Ysp2, a cryptococcal sterol transporter of the LAM family, impacts the ability of cryptococci to maintain key cellular structures, survive in the mammalian host, and cause disease. We found that *ysp2*Δ cells grown in host-like conditions present with striking invaginations of both the PM and the cell wall, as well as increased levels of ergosterol at the PM (Fig 4).

At the ER, WT *C. neoformans* cells synthesize ergosterol, which is then transported to the PM by a yet-to-be identified anterograde sterol transporter. We propose that Ysp2 functions as a retrograde transporter, removing excess ergosterol from the PM to balance the system. In the host, there is likely an increased need for efficient ergosterol redistribution, to rapidly respond to environmental stress. Based on our observations, we hypothesize that when Ysp2 is absent in this situation, retrograde transport of sterol is abrogated while anterograde transport continues (Fig. 8A), leading to an accumulation of ergosterol at the PM (Fig. 8B). The excess ergosterol causes the PM to invaginate (Fig. 8C), likely by increasing membrane fluidity (40). This in turn causes mislocalization of PM-resident proteins, including proteins involved in cell wall synthesis, and results in enriched synthesis of new cell wall material (darker green in Fig. 8D) within the invaginated areas.

**Fig 8.**
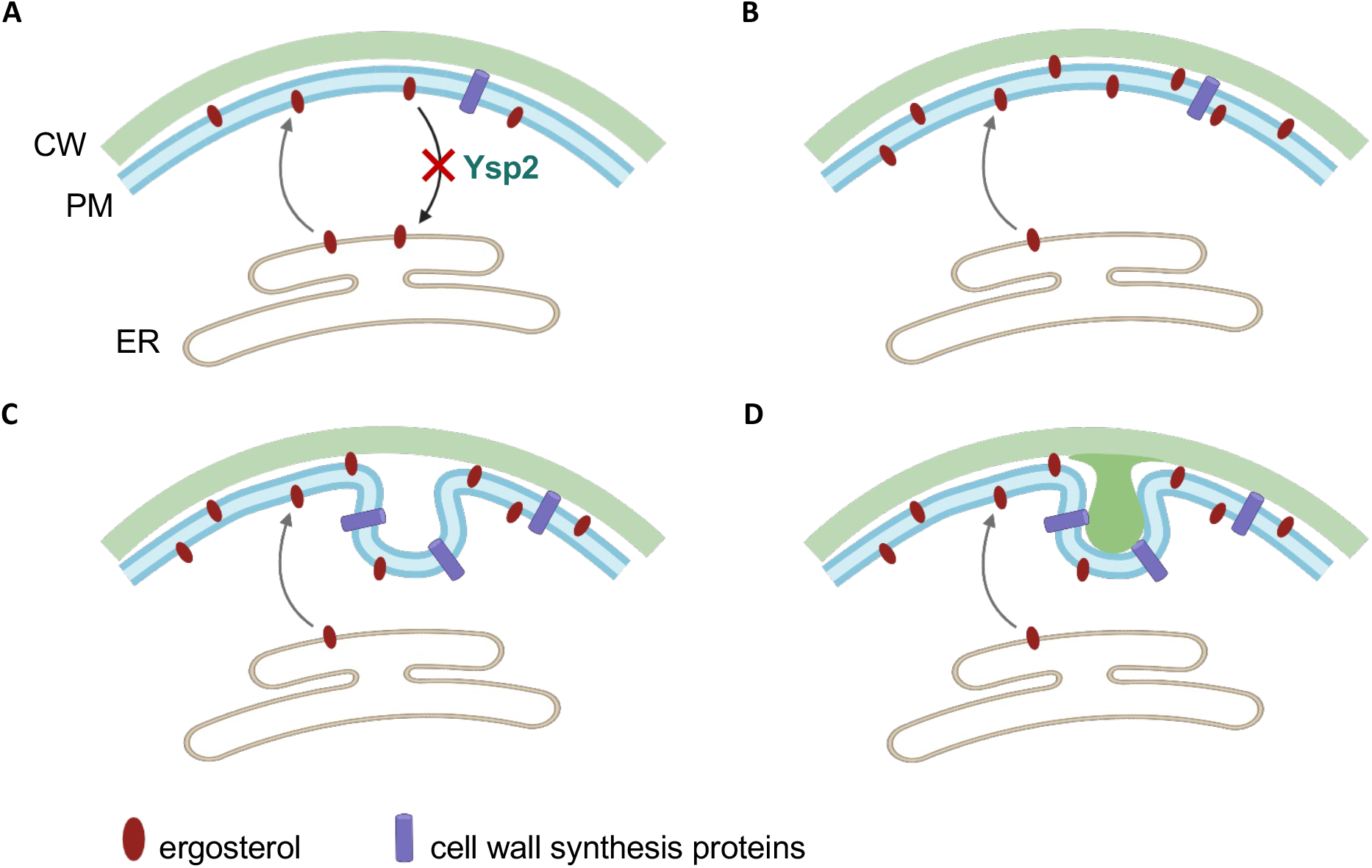
Model of Ysp2 function and how its absence influences plasma membrane and cell wall morphology. CW, cell wall; PM, plasma membrane; ER, endoplasmic reticulum.

Our evidence for protein mislocalization comes in part from studies of Pma1, a PM ATPase that is associated with lipid rafts (41, 42, 31). Lipid rafts require tight associations of sterols with sphingolipids <u>(1, 2)</u>; we speculate that ergosterol accumulation at the PM perturbs these structures, resulting in aberrant protein localization (43–45). In support of this idea, Pma1 in the *ysp2*Δ mutant appeared as puncta at the cell surface and occasionally in vacuoles (Fig 5A and S4A), in contrast to its smooth PM distribution in WT cells (46). We expect that changes in membrane organization also affect the localization of other PM-resident proteins, such as phospholipase B1 (Plb1), a virulence factor (47), and multiple cell wall synthesis proteins (e.g. Chs3, Cda1, Cda2, and Fks1 (48–51)). The latter group is particularly critical during cell growth, when fungi actively remodel and thicken their cell walls in response to changes in their environment (52–54). We hypothesize that mislocalization of such proteins leads to the aberrant wall synthesis suggested in our model (Fig. 8).

In addition to cell wall alterations, we noted altered capsule architecture in *ysp2*Δ cells, which exhibited both reduced capsule thickness and increased capsule permeability. These phenotypic alterations may be due to changes in two compartments that are impacted by sterol organization: the cell wall and the ER. The protein mislocalization that occurs in *ysp2*Δ cells, mentioned above, likely changes cell wall composition and structure, an idea supported by the changes in chitin revealed by our CFW staining (Fig 4D). The combination of PM and wall changes may further lead to the mislocalization of capsule attachment and remodeling proteins (e.g. Ags1, Pbx1, or Lhc1) and/or altered ability to attach capsule fibers (52, 53, 55–57).

Other potential explanations for the capsule alterations observed in *ysp2*Δ cells relate to ER function. Because this organelle is the site of ergosterol synthesis and esterification, loss of retrograde ergosterol transfer will indirectly change ER ergosterol levels, potentially influencing secretory processes and thus the export of capsular polysaccharides. This idea is supported by a previous study, which showed reduced secretion of capsule polysaccharides in a mutant defective in ER homeostasis (58). Changes in ER lipid composition could also alter the localization or activity of ER-resident nucleotide sugar transporters such as Uut1 and Uxt2, which are required for capsule synthesis (59, 60). Notably, proteins involved in ergosterol synthesis are not strongly implicated in capsule formation: the ergosterol synthesis mutant *erg6*Δ has normal capsule, although the synthesis regulator mutant *sre1*Δ does show slight capsule defects (33, 36, 37, 39). Overall, our results suggest a connection between ergosterol organization and capsule elaboration, which warrants further exploration.

High levels of ergosterol at the PM could potentially be explained by increased synthesis of this compound or aberrant distribution of normal amounts. Based on the putative role of Ysp2 as a sterol transporter, we initially assumed that the latter explained the phenotypes we observed. Consistent with this idea, even though our microscopy and flow analyses showed increased ergosterol at the cell surface, sterol quantification indicated no changes in total abundance (Fig 6A). Nonetheless, our qPCR data suggests a more complex story, since *ysp2*Δ cells grown in a host-like environment showed consistent 2-3-fold upregulation of multiple genes involved in ergosterol synthesis (Fig 6B). We speculate that this stimulation is triggered by the relative lack of ergosterol *within* the cell, which occurs because mutant cells cannot retrieve it from the PM. Precedent for this idea comes from studies of the *S. cerevisiae* sterol-regulatory element-binding protein Upc2, which is activated by depletion of intracellular ergosterol (6). We suggest a similar mechanism for the cryptococcal sterol regulator, Sre1 (36). The resulting increased ergosterol synthesis, although it represents an effort by the cell to maintain homeostasis, may in fact contribute to the skewed sterol distribution caused by reduced retrograde transport.

The discussion above still leaves open the question of why we see no changes in total ergosterol in *ysp2*Δ cells if synthesis is being increased, albeit modestly. The dramatic increase in lipid droplets that we observe in the mutant (Fig 6C) suggests an answer. In *S. cerevisiae*, excess sterols are not degraded; instead, they are either esterified and stored in lipid droplets, or secreted into the environment as sterol acetates (6). If similar events occur in *C. neoformans*, the increase in lipid droplets likely reflects the esterification and storage of sterols made in response to upregulation of the ergosterol synthesis pathway. We did not measure esterified sterols, so they are not included in the quantification of free ergosterol, but future experiments to directly quantify these and other related compounds could address this model (Fig 6C). This hypothesis could also be tested by looking for potential upregulation of Are1 and Are2, ER-resident enzymes that esterify sterols (6).

Multiple factors likely contribute to the reduced virulence of *ysp2*Δ. One is the poor growth of this mutant in host-like conditions (Fig 2D, S2). Its inability to survive within host phagocytes may also reduce virulence by impeding cryptococcal dissemination in the host (24). The changes that occur in both the capsule and cell wall may further contribute to the reduced virulence, as these are well-characterized virulence factors with critical roles in protecting fungal cells from external stresses (26). A last possibility is that accumulation of ergosterol at the mutant PM affects the host immune response, as work in several fungal pathogens has shown that the level of cell surface ergosterol correlates with the ability of fungal pathogens to trigger host pyroptosis (61, 62).

We have focused on the *C. neoformans* homolog of the *S. cerevisiae* LAM family protein Ysp2. Curiously, *C. neoformans* encodes only one LAM homolog, although *S. cerevisiae* has six such proteins with overlapping activities (Fig 1A) (19, 63). It may be that cryptococcal Ysp2 serves additional functions compared to the *S. cerevisiae* protein. This idea is supported by the distinct localization of Ysp2 in the two organisms. In the model yeast Ysp2 is an ER-resident protein that localizes to ER-PM contact sites at the cell periphery (19, 21), while other LAM family members act at ER-PM, ER-mitochondria, or ER-vacuole contact sites (19, 20). In contrast, we observed both intracellular and peripheral localization of *C. neoformans* Ysp2. This suggests that it acts at additional membrane interfaces, such as those of mitochondria or vacuoles, as well as at surface contact sites. Another possible explanation for the single LAM family protein in *C. neoformans* is that additional proteins involved in *C. neoformans* sterol transport, which lack homology to this family yet perform similar functions, remain to be discovered.

We have identified a retrograde sterol transporter in *C. neoformans*, Ysp2, that is critical for virulence by influencing key cellular functions including PM integrity, cell wall formation, capsule elaboration, and lipid homeostasis. We provide a model that explains the phenotypes observed in *ysp2*Δ and predicts how cells respond to excess ergosterol accumulation. Our findings provide insights into the role of sterol transport in cryptococcal biology, particularly in the context of the host. Beyond the future directions mentioned above, these discoveries suggest multiple important topics that remain to be explored, including the complete set of proteins responsible for sterol transport in *C. neoformans*, how sterol accumulation changes the biophysical properties of the cryptococcal PM, and ergosterol regulation and homeostasis in the environment of the infected host.

## METHODS

### Cell growth and strain construction

*C. neoformans* strains were grown overnight in YPD medium (1% [wt/vol] Bacto yeast extract, 2% [wt/vol] dextrose, 2% [wt/vol] Bacto peptone in double-distilled water [ddH_2_O]) at 30°C with shaking at 230 rpm, collected by centrifugation, washed twice with sterile PBS, diluted to 10^6^ cells/mL in RPMI and incubated at 37°C in 5% CO_2_ for 24 hours in 6-well plates or T-75 tissue culture flasks. For growth curves, cells were grown overnight in YPD, washed, and adjusted to 1 × 10^5^ cells/ml for growth in YPD (30°C, 37°C, or 37°C + 5% CO_2_), RPMI at 37°C + 5% CO_2_ (37R5), or DMEM at 37°C + 5% CO_2_ (37D5).

Strain construction is detailed in Text S1.

### Virulence studies

All animal protocols were approved by the Washington University Institutional Animal Care and Use Committee (Protocol #20-0108), and care was taken to minimize animal handling and discomfort.

For survival studies, groups of ten 8-week-old female C57BL/6 mice (The Jackson Laboratory) were anesthetized by injection of 1.20 mg ketamine and 0.24 mg xylazine in 110 μl sterile PBS and intranasally infected with 1.25 × 10^4^ cryptococcal cells. The mice were monitored and humanely sacrificed when their weight decreased to below 80% of their initial weight or if they showed signs of disease. To assess organ burden at the time of sacrifice the lungs and brains were harvested, homogenized, diluted, and plated on YPD agar. The resulting CFUs were enumerated, and survival differences were assessed by Kaplan-Meier analysis.

### Fungal intracellular survival

We obtained bone marrow-derived macrophages (BMDM) as detailed in Text S1. Fungal cells from overnight YPD cultures were washed in PBS, opsonized in 20% human serum in PBS (10^7^ cells/ml, 30 min, 37°C), washed in PBS, and resuspended in RPMI. Opsonized fungi were added to BMDM cells at an MOI of 0.1 and incubated for 1.5 hours to permit engulfment. Wells were then washed and refilled with prewarmed BMDM medium and plates were incubated for 0, 24, or 48 hours at 37°C, 5% CO_2_; washed twice with sterile PBS; refilled with sterile ddH_2_O; incubated at room temperature for 30 minutes to lyse BMDM; and plated on YPD agar to quantify CFU.

### Microscopy and flow cytometry

For imaging, fungal strains were grown as above but resuspended at 10^7^ cells/ml for staining as detailed in Text S1, and then imaged using a ZEISS Axio Imager M2 fluorescence microscope or a ZEISS LSM880 confocal laser scanning microscope. Electron microscopy was performed as detailed in Text S1. For flow cytometry, cells were resuspended in 1 ml of PBS with 10 mM NaN_3_. Data were acquired on a BD LSRFortessa Cell Analyzer and analyzed using FlowJo software.

### Phenotyping

Cells grown as above were adjusted to 10^7^ cells/ml in PBS and serially diluted to final cell concentrations of 10^6^, 10^5^, 10^4^, and 10^3^ cells/ml. Four microliters of each dilution were spotted onto YPD and stress plates and grown at 30 and 37°C. To impose membrane stress, YPD agar was supplemented with 0.01% SDS, 1.2 M NaCl, 1 µg/ml amphotericin B, and 8 µg/ml fluconazole. YPD agar was supplemented with 0.2% calcofluor white (wt/vol) and 0.05% Congo red for cell wall stress, or 0.125 µg/ml tunicamycin for ER stress.

### Sterol Analysis

Lipid extraction was performed as previously described (64). For details of lipid extraction and GC-MS analysis, see Text S1.

### qPCR

Total RNA was extracted using TRI-Reagent (Applied Biosystem) and cDNAs were synthesized using the SuperScript III First-Strand Synthesis System SuperMix kit (Invitrogen) for quantitative PCR analysis using the SYBR® Green PCR Master Mix kit (Applied Biosystems) as recommended by the supplier and a CFX96 Touch Real-Time PCR Detection System (Bio-Rad). Relative gene expression was calculated using the CT comparative method (2^−ΔΔCT^), with *ACT1* expression as a normalization control.

## Supporting information

Choy et al Supplementary Methods

Choy et al Supplementary Figures

## ACKNOWLEDGMENTS

We thank Liza Loza for assistance with animal experiments, Wandy Beatty of the Molecular Microbiology Imaging Facility for expert electron microscopy, Cheryl Frankfater of the Biomedical Mass Spectrometry Resource at Washington University in St. Louis for GC-MS analysis of sterols, Mark Bradley for amphotericin B-Cy5, and Tom Kozel for mAb 302. We are grateful to the members of the Doering lab and Andrew Jezewski for helpful discussions and to Daphne Ko, Liza Loza, and Thomas Hurtaux for comments on the manuscript.

This work was supported by NIH grants R21 AI136688, R21 AI140979, and R01 AI135012 to T.L.D. H.L.C and E.A.G. were also partly supported by T32 GM007067, and E.A.G by a Sondra Schlesinger Graduate Fellowship from the Department of Molecular Microbiology, Washington University School of Medicine.

## AUTHOR CONTRIBUTIONS

H.L.C: Conceptualization, Methodology, Investigation, Validation, Writing – Original Draft, Writing – Review & Editing. E.A.G: Investigation, Validation, Writing – Review & Editing. T.L.D: Conceptualization, Writing – Original Draft, Writing – Review & Editing, Supervision, Project Administration, Funding Acquisition.

