## Supplementary material for "Ergosterol distribution controls surface structure formation and fungal pathogenicity": Choy et al Supplementary Methods

### Supplementary Text S1

#### Strain construction

All *C. neoformans* strains used were in the serotype A KN99α background. For strain construction, we used a split-marker strategy (1) with biolistic transformation, selected candidates by patterns of drug resistance, and confirmed all candidate transformants by PCR and whole genome sequencing. We first generated two *ysp2*Δ deletion strains by replacing *YSP2* with either a G418 (2) or NAT resistance marker (3). To complement these mutants at the native site, we replaced the G418 or NAT coding sequence with the *YSP2* coding sequence, preceded by a NAT or G418 resistance marker, respectively. To endogenously tag *YSP2*, we used a tagging module consisting of codon-optimized mNeonGreen, a tandem Calmodulin Binding Peptide-2X FLAG Tag, and a NAT resistance marker.

We used a CRISPR-Cas9 strategy to tag the C-terminus of Pma1 (*CNAG_06400*) with an mNeonGreen tagging module as described above in tandem with a G418 marker (4).

#### BMDM Preparation

For bone marrow-derived macrophages (BMDM), we isolated bone marrow from C57BL/6 mouse femurs and expanded the cellular population by growth in BMDM medium (RPMI supplemented with 10% heat-inactivated FBS and 20% L cell supernatant) for seven days. Macrophages were isolated using anti-F4/80 conjugated biotin and anti-biotin magnetic microbeads.

#### Microscopy

For imaging, fungal strains were grown as in the main text and resuspended at $10^7$ cells/ml for staining with Lucifer Yellow (200 µg/ml, 30 min, RT), calcofluor white, (100 µg/ml, 15 min, RT), filipin (5 µg/ml 10 min, RT), amphotericin B-Cy5 (10 µM, 45 min, 37˚C), or Nile Red (0.0005%, 15 min, RT) (5). For capsule imaging, cells were resuspended in 600 µl PBS and mixed with 300 µl India Ink or stained (30 min, RT) with 50 µg/ml of anticapsular monoclonal antibody 302 conjugated to Alexa Fluor 488 (Molecular Probes).

For electron microscopy, cells were grown as described above and fixed in 2.5% glutaraldehyde (Ted Pella Inc., Redding, CA) in 0.1 M sodium cacodylate buffer for 1 h at room temperature and then overnight at 4°C. Samples were washed in sodium cacodylate buffer and postfixed in 1% osmium tetroxide (Ted Pella Inc.) for 1 h at room temperature before dehydration by successive 30 min incubations in water–ethanol mixtures with increasing ethanol concentrations. The ethanol-substituted samples were then substituted in propylene oxide (twice for 30 min each) and infiltrated and embedded in Eponate 12 resin (Ted Pella, Redding, CA). Blocks were polymerized overnight at 65°C. Ultrathin sections of 95 nm were cut with a Leica Ultracut UCT ultramicrotome (Leica Microsystems, Bannockburn, IL), stained with uranyl acetate and lead citrate, and viewed on a JEOL 1200 EX transmission electron microscope (JEOL USA Inc., Peabody, MA) equipped with an AMT 8-megapixel digital camera and AMT Image Capture Engine V602 software (Advanced Microscopy Techniques, Woburn, MA).

### Sterol Analysis

Lipid extraction was performed as previously described (6). Briefly, 2.5 x $10^8$ cells were washed twice in sterile water and resuspended in 1.5 ml of Mandala lipid extraction buffer (ethanol:d$H_2O$:diethylether:pyridine:14.2 N ammonium hydroxide; 15:15:5:1:0.018; v:v:v:v:v) with 5 µg of cholesterol added as an internal standard (7). Samples were vortexed, sonicated (Fisher Scientific Sonic Dismembrator Model 300; 40% power for 30 s) three times at intervals of 30 s, and incubated at 60˚C for 15 min. This process was repeated, cell debris was removed by centrifugation, the supernatant fraction was evaporated, and the residue was resuspended in chloroform:methanol (1:2; v:v) and incubated at 37˚C for 1 h. 1 ml chloroform and 1 ml d$H_2O$ were added to initiate phase separation and the hydrophobic layer was removed, dried, and subjected to mild alkaline hydrolysis (1 h, RT) by the addition of 0.5 ml chloroform and 0.5 ml 0.6 M methanolic KOH. 0.325 ml 1M HCl and 0.125 ml d$H_2O$ were then added for phase separation, and the hydrophobic layer was dried and stored at -20˚C until analyzed.

Gas chromatography and mass spectrometry analyses were performed at the Biomedical Mass Spectrometry Core. The dried lipid residue was resuspended in 100 µl of a mixture of 1:0.8:2.2 (v:v:v) BSTFA with 1% TMCS:pyridine:acetonitrile for derivatization, vortexed, heated at 65°C for 1 h, and vortexed again. Electron ionization (EI) GC-MS analysis was conducted on an Agilent (Santa Clara, CA USA) 7890A GC coupled with Agilent 5975C MSD using a 25-m Agilent J & W capillary column (DB-1; inner diameter, 0.25 mm; film thickness, 0.1 µm). An aliquot of 2 µl of the derivatized solution was injected with a 10:1 split onto the GC/MS system controlled by Agilent ChemStation software. The temperature program started at 85 °C and held for 1 min, ramped 50°C/min to 268°C, held at 268°C for 1 min, increased 1°C/min to 275°C, increased 5°C/min to 282°C, and increased 20°C/min to 300°C with a final hold at 300°C for 11.1 min. Mass spectra were collected in full scan mode from m/z 60 to 525 and sterol peaks were identified by matching the built-in NIST14 library. Endogenous ergosterol and the added cholesterol internal standard peaks were integrated and quantified using a standard curve established by a series of injections of various ergosterol/cholesterol ratio mixtures.
