## Supplementary material for "Ergosterol distribution controls surface structure formation and fungal pathogenicity": Choy et al Supplementary Figures

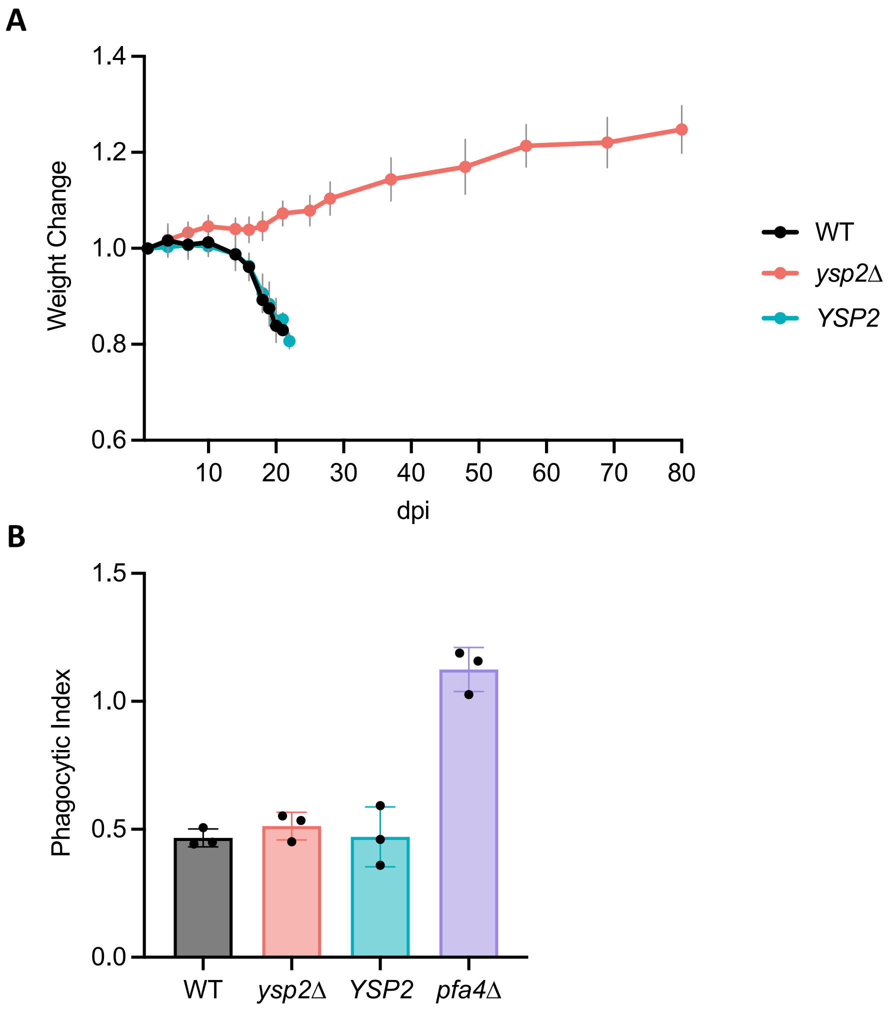


**Fig S1.** Virulence and viability of *ysp2*Δ cells. (A) Weight of mice infected with the indicated strains over time, normalized to initial weight. Mean ± SD are displayed for 10 mice per group. (B) Phagocytic index (internalized fungi/host cells) of the indicated strains after 1 h incubation as described in the Methods. *pfa4*Δ, positive control strain for increased phagocytic index (43). Mean ± SD of three independent experiments is shown. (C) The indicated strains were grown in 37R5 and sampled at the times shown for their ability to form colonies on solid YPD medium. Mean ± SD are displayed for technical triplicates of a typical experiment.

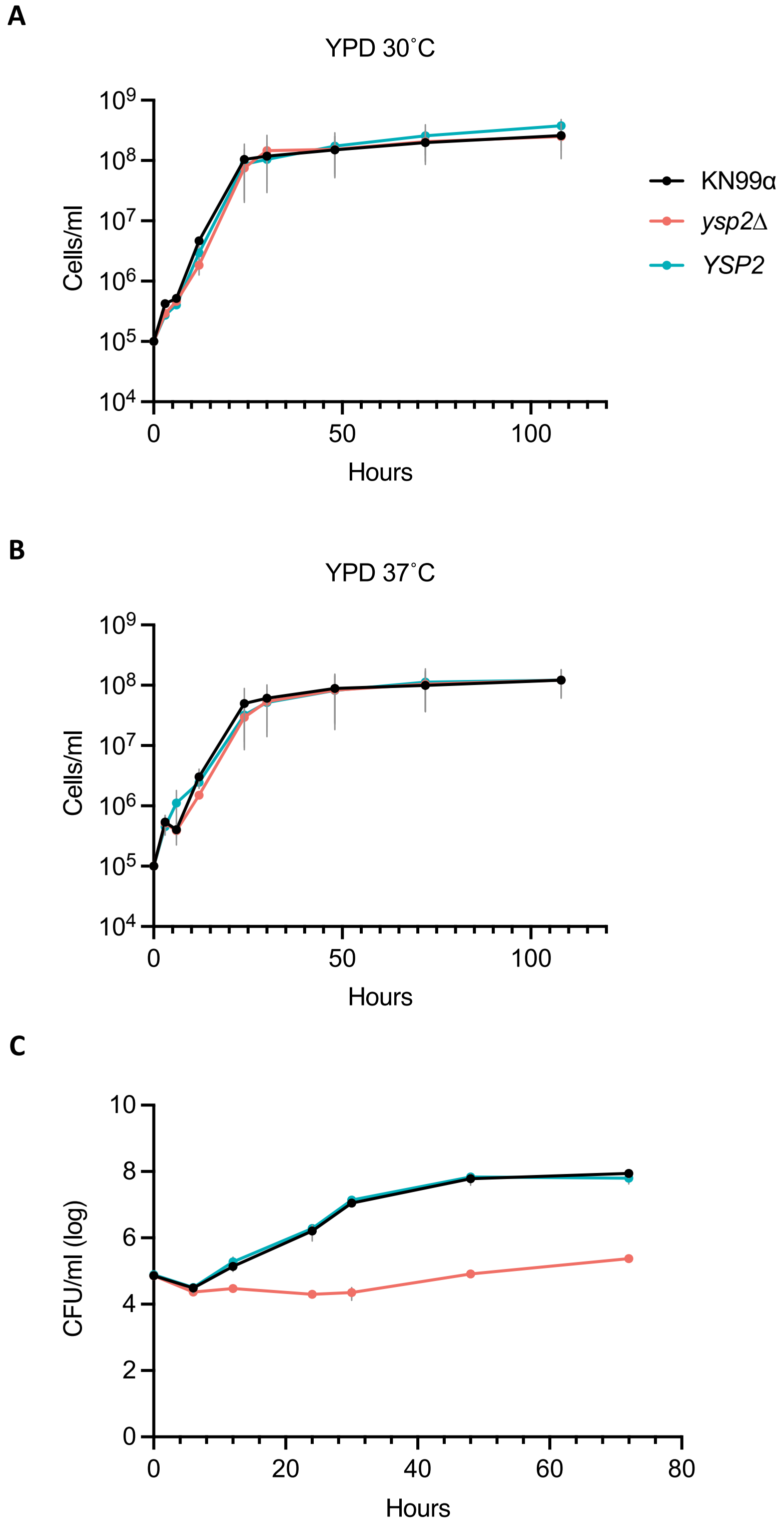


**Fig S2.** Growth curves in the conditions shown (mean ± SEM of three independent experiments).

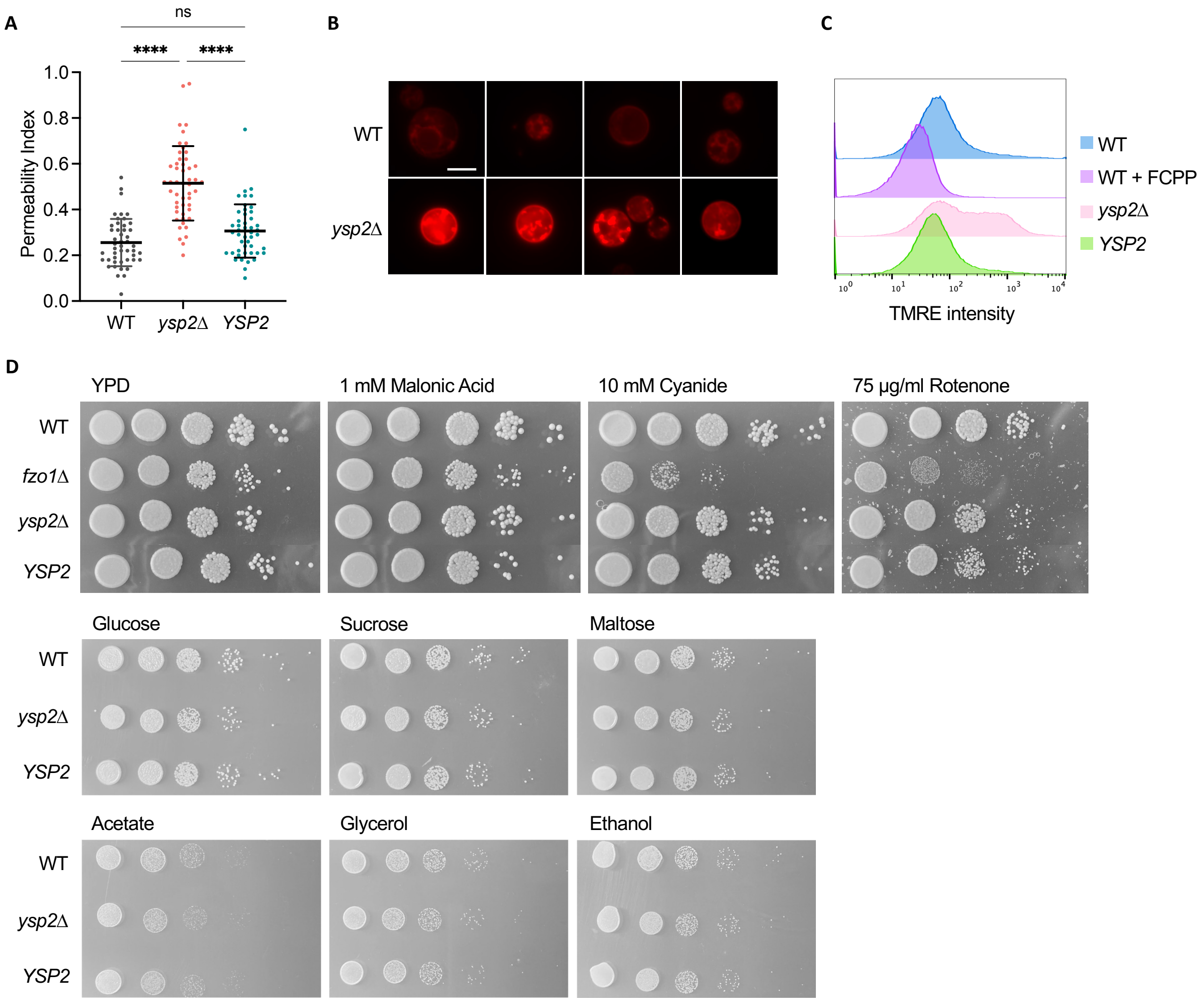


**Fig S3.** Capsule permeability and mitochondrial functions. (A) Capsule permeability of the indicated strains measured as the fraction of the capsule radius (defined by India ink exclusion) that is penetrated by 2,000 kDa Dextran beads. >40 cells were quantified per sample, plotted with mean ± SD . (B) Representative images of cells grown in 37D5 + 1 mM $H_2O_2$ stained with 1 mM MitoTracker CMXRos. All images are captured with the same parameters and shown to the same scale; bar, 5 μm. (C) TMRE fluorescence intensity quantified by flow cytometry of the indicated strains after 24 h growth in YPD. FCPP, carbonyl cyanide 4-(trifluoromethoxy) phenylhydrazone, an electron transport chain uncoupler. (D) Stress phenotypes of the indicated strains on rich medium (YPD). Serial dilutions of the indicated strains were grown in the absence or presence of the indicated additives. Alternative carbon sources were tested at 2% (wt/vol).

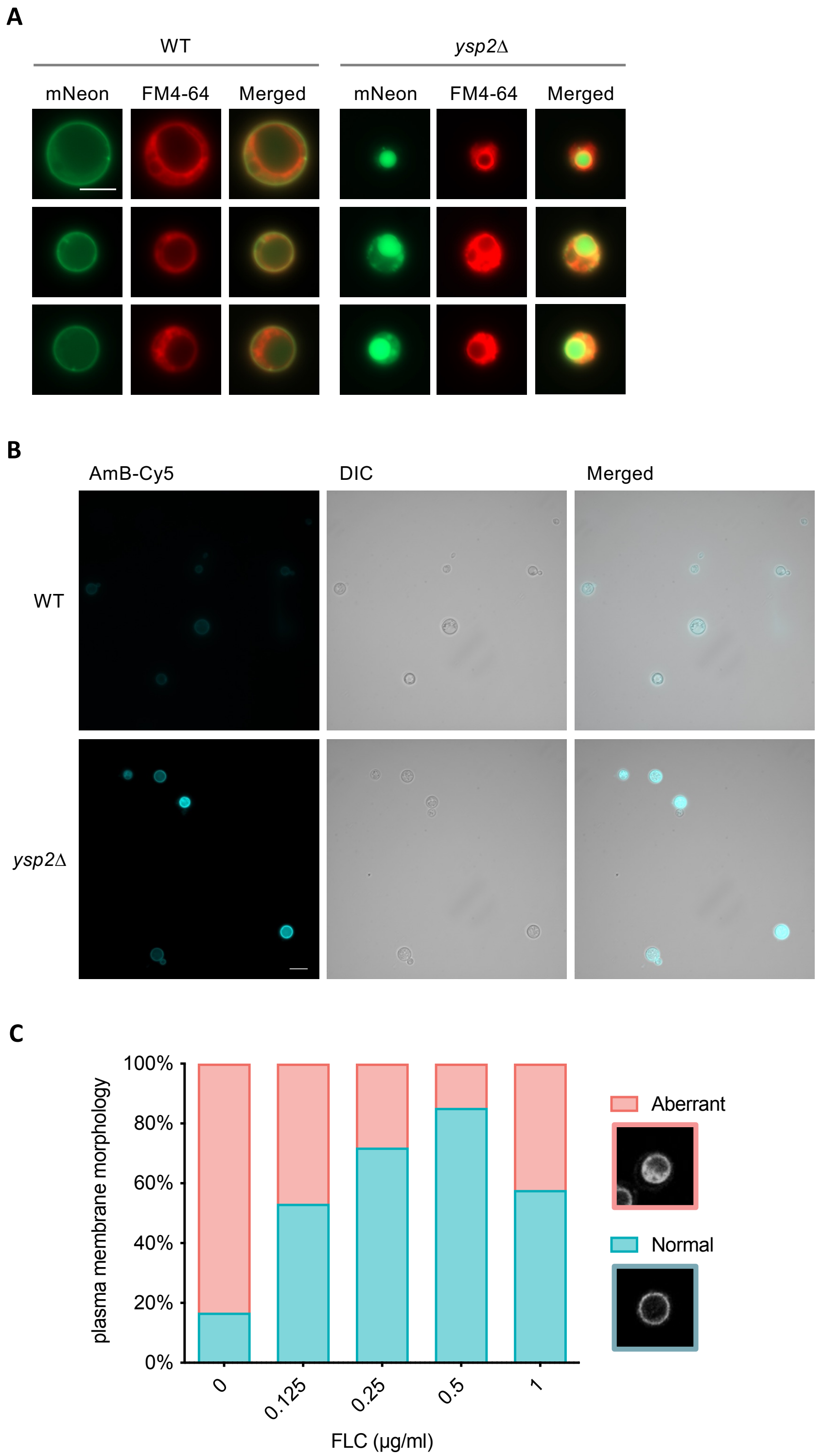


**Fig S4.** Capsule permeability and mitochondrial functions. (A) Representative fluorescence micrographs of WT and mutant strains with mNeonGreen-tagged Pma1 and the vacuole counterstained with FM4-64. All images are to the same scale; bar, 5 μm. (B) Representative fluorescence images of cells stained with 10 μM AmB-Cy5. All images are to the same scale; bar, 10 μm. (C) Rescue of *ysp2*Δ plasma membrane invagination by fluconazole. Cells were grown in 37R5 and the fraction of aberrant cell membranes quantified at 24 h. 30 cells were scored per condition.
